# Newly raised anti-c-Kit antibody detects interstitial cells of Cajal in the gut of chicken embryos

**DOI:** 10.1101/2022.05.26.493534

**Authors:** Rei Yagasaki, Yuuki Shikaya, Teruaki Kawachi, Masafumi Inaba, Yuta Takase, Yoshiko Takahashi

## Abstract

The gut peristaltic movement, a wave-like propagation of a local contraction, is important for the transportation and digestion of ingested materials. Among three types of cells, the enteric nervous system (ENS), smooth muscle cells, and interstitial cells of Cajal (ICCs), the ICCs have been thought to act as a pacemaker, and therefore it is important to decipher the cellular functions of ICCs for the understanding of gut peristalsis. c-Kit, a tyrosine kinase receptor, has widely been used as a marker for ICCs. Most studies with ICCs have been conducted in mammals using commercially available anti-c-Kit antibody. Recently, the chicken embryonic gut has emerged as a powerful model to study the gut peristalsis. However, since the anti-c-Kit antibody for mammals does not work for chickens, cellular mechanisms by which ICCs are regulated have largely been unexplored. Here, we report a newly raised polyclonal antibody against the chicken c-Kit protein. The specificity of the antibody was validated by both Western blotting analyses and immunocytochemistry. Co-immunostaining with the new antibody and anti-α smooth muscle actin (αSMA) antibody successfully visualized ICCs in the chicken developing hindgut in the circular muscle- and longitudinal muscle layers: as previously shown in mice, common progenitors of ICCs and smooth muscle cells at early stages were double positive for αSMA and c-Kit, and at later stages, differentiated ICCs and smooth muscle cells exhibited only c-Kit and αSMA, respectively. A novel ICC population was also found that radially extended from the submucosal layer to circular muscle layer. Furthermore, the new antibody delineated individual ICCs in a cleared hindgut. The antibody newly developed in this study will facilitate the study of peristaltic movement in chicken embryos.

## Introduction

Gut peristaltic movements, a wave-like propagation of a local contraction, play important roles in the effective transportation and digestion/absorption of ingested materials. Although macroscopic physiology of gut peristalsis has extensively been studied in adults, the cellular mechanisms underlying the peristaltic regulation remain largely unexplored. Three types of cells play important roles in the peristaltic regulations: enteric nervous system (ENS), smooth muscle cells, and interstitial cells of Cajal (ICCs). While ENS is derived from neural crest cells, both the smooth muscle cells and ICCs originate from common progenitors of splanchnopleural (splanchnic) mesoderm (Lecoin et al., 1996; Torihashi et al., 1997). In mice, it has been proposed that ICCs act as a pacemaker for the peristaltic movement, since knockout mice of the *c-Kit* gene, a tyrosine kinase receptor expressed in ICCs, exhibited severe defects in both ICC differentiation and peristaltic rhythm(Der-Silaphet et al., 1998; Ordög et al., 2002).

It is known in mice that common progenitors of ICCs and smooth muscle cells express c-Kit and α smooth muscle actin (αSMA) at early stages (Midrio et al., 2004; Radenkovic et al., 2010). Subsequently, only cells that receive stem cell factor (SCF; the c-Kit ligand) differentiate into ICCs. At later stages, differentiated ICCs can be identified as c-Kit-positive and aSMA-negative, whereas smooth muscles cells are αSMA-positive and c-Kit-negative (Klüppel et al., 1998; Radenkovic et al., 2010). This also highlights a usefulness of anti-c-Kit antibody, and indeed in mammals, commercially available antibodies have widely been used to study ICCs in embryos and adults (Baker et al., 2021; Maeda et al., 1992; Malysz et al., 2017; Mei et al., 2009; Roberts et al., 2010; Torihashi et al., 1999; Torihashi et al., 1995; Torihashi et al., 1997; Ward et al., 1994; Wester et al., 1999).

The gut musculature is composed of longitudinal muscles (LM) and circular muscles (CM). In mice, cells called myenteric ICCs (ICC-MY) residing in between LM and CM exhibit multiple cellular processes and form a cellular network (Iino et al., 2011; Iino et al., 2007; Nemeth & Puri, 2001). ICCs localized within LM and CM layers are called intramuscular ICCs (ICC-IM), which are bipolar in shape and aligned with elongated muscle cells. It has been proposed that ICC-MY acts as a pacemaker, whereas ICC-IM plays a role in signal transduction within networks (Burns et al., 1996; Huizinga et al., 1995; Ward et al., 1994). However, many of these studies have been performed with cultured ICCs or gut fragments, and therefore how these ICCs contribute to the peristaltic regulation *in vivo* remains elusive.

The chicken embryonic gut, which is highly gene-manipulable *in/ex vivo*, has recently emerged as a powerful model for elucidating the endogenous potential and regulation of peristalsis-related cells including ICCs, because peristalsis occurs even though no ingested material is present in the moving gut (Chevalier et al., 2017). In addition, the distribution map of peristaltic movements along the entire gut axis is available (Shikaya et al., 2022). Previous studies using *in situ* hybridization with *c-Kit* showed that the composition of ICCs in the chicken gut largely parallel that in mammals (Chevalier et al., 2020; Lecoin et al., 1996; Yang et al., 2012). For example, an ICC network-like structure between the LM and CM (probably ICC-MY) was reported (Chevalier et al., 2020), and ICC-MY differentiates at the time of LM differentiation (Lecoin et al., 1996). However, unlike mammals, the antibody that detects the chicken c-Kit protein was not available, and this delayed the investigation at the cellular level with ICCs in the chicken embryonic gut.

In this study, we report a newly raised antibody against the chicken c-Kit protein. The specificity of the anti-serum/polyclonal antibody to the chicken c-Kit protein was validated by both Western blotting analysis and immunocytochemistry with c-Kit cDNA-transfected cells. Furthermore, the newly raised anti-c-Kit antibody visualized c-Kit-positive ICC populations in histological sections, and it also delineated individual cells of ICC-MY and ICC-IM in a cleared hindgut. The antibody obtained in this study will facilitate the study of ICCs and gut peristalsis in chickens.

## Materials and Methods

### Polypeptides for rabbit immunization

The polypeptide at the position of 716 to 728 amino acids (aa) of the chicken c-Kit protein (Fig. 1A) was synthesized and used for immunization of a rabbit. The antiserum was harvested, and purified by affinity purification using the aforementioned polypeptide (Sigma Aldrich Japan).

**Figure 1.**
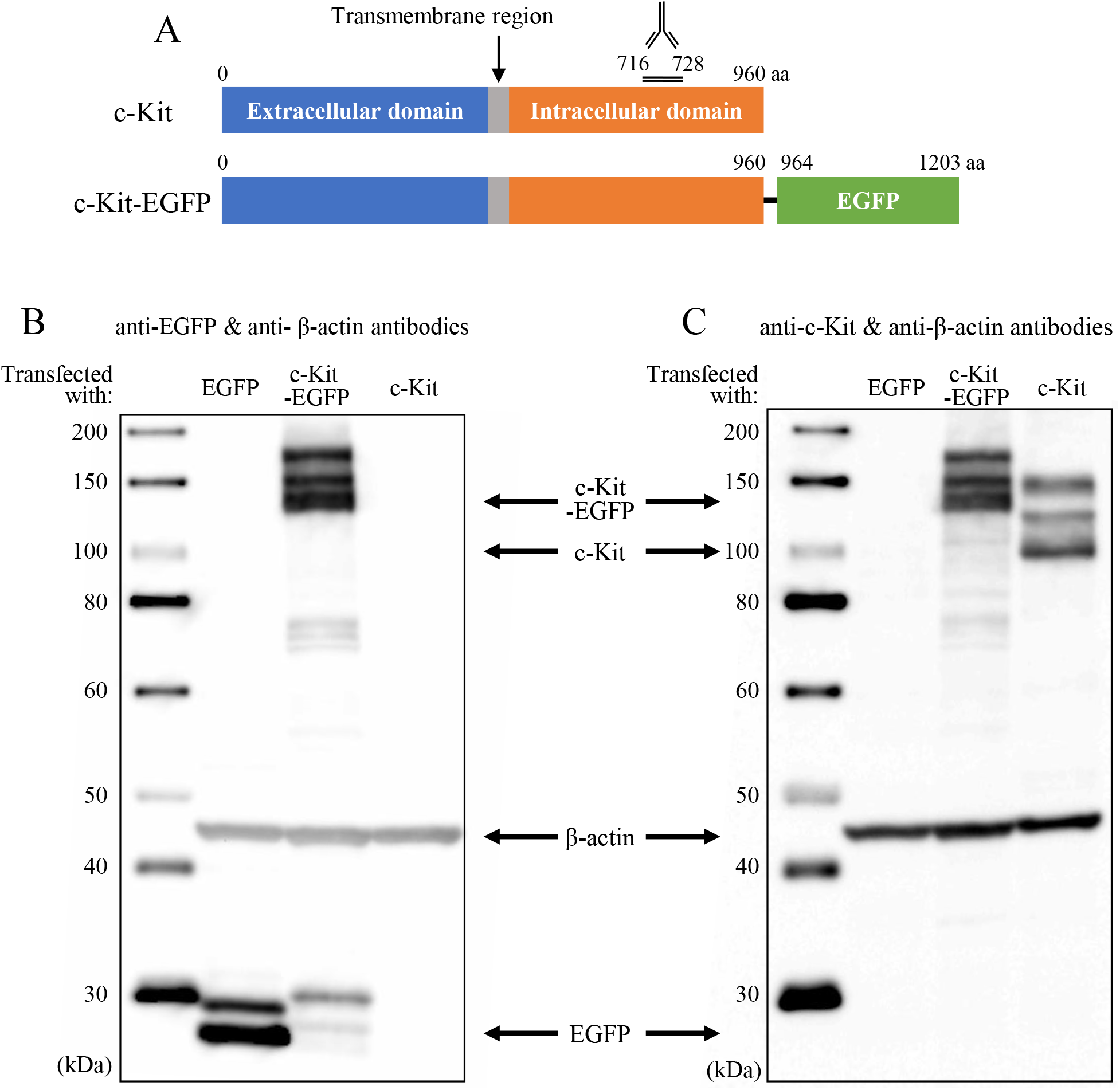
(A) Polypeptide synthesized at the position of 716 to 728 of the chicken c-Kit protein was used for immunization of a rabbit. (B, C) Obtained polyclonal antibody was validated by Western blotting analyses using lysates of DF1 cells transfected with *c-Kit, c-Kit-EGFP* (fusion) or *EGFP* cDNAs. The left lane of each panel shows size markers. Signals were detected with (B) 180, (C) 270-second exposure.

### Chicken embryos

Fertilized chicken eggs were obtained from the Yamagishi poultry farms (Wakayama, Japan). Embryos were staged according to embryonic days. All animal experiments were conducted with the ethical approval of Kyoto University (#202110).

### Plasmids

Full-length cDNA of chicken *c-Kit* (GeneBank: D13225.1, CDS: 2883 bp) was PCR-amplified: forward 5’-GGGGCGGCCGCGCCACCATGGAGGGCGCGCACCTCGC-3’, reverse 5’-GGGGCGGCCGCTCAAACATCTTCGCGTACCAGGA-3’. pT2AL200R175-CAGGS-LynEGFP-IRES-Neo_r_ (Inaba et al., 2019) was digested with NotI to remove LynEGFP, into which DNA fragments were inserted at the NotI site to produce: pT2AL200R175-CAGGS-c-Kit-IRES-Neo_r_, pT2AL200R175-CAGGS-c-Kit-EGFP-IRES-Neo_r_

c-Kit-EGFP fusion protein: chicken c-Kit cDNA was inserted into multiple cloning site of pEGFP-N1 Vector (Clontech, #6085-1) including EGFP sequences using a primer set: forward 5’-CCACCGGTCGCCACCATGGAGGGCGCGCACCTCGC-3’, reverse 5’-GCTCACCATCTCGCGTCCAACATCTTCGCGTACCA-3’. Subsequently, cDNA of *c-Kit-EGFP* was generated by PCR using the following primers: forward 5’-AGAATTCCTCGAGCGGCCGCATGGAGGGCGCGCACCTCGC-3’, reverse: 5’-TGGACGAGCTGTACAAGTAAGCGGCCGCCACCGCGGTGGA-3’.

### DNA transfection into cultured cells

Transfection of DNA plasmids into the chicken fibroblast line cells DF1 was performed using Lipofectamin 3000 (Invitrogen). Three kinds of transfected cells were separately prepared introduced with: pT2AL200R175-CAGGS-EGFP-IRES-Neo^r^, pT2AL200R175-CAGGS-c-Kit-EGFP-IRES-Neo^r^, and pT2AL200R175-CAGGS-c-Kit-IRES-Neo^r^. After 48 hours of transfection, these cells were harvested.

### Western blotting

For Western blotting analyses, transfected DF1 cells were homogenized in phosphate buffer saline (PBS: 0.14 M NaCl, 2.7 mM KCl, 10 mM Na_2_HPO_4_-12H_2_O, 1.8 mM KH_2_PO_4_). The whole cell lysate was mixed with a Laemmli sample buffer (BIORAD, 161-0737) and heated for 5 min at 95 °C. The eluates were subjected to sodium dodecyl sulfate-polyacrylamide gel electrophoresis (SDS-PAGE) with a marker (SMOBIO, WM1000), followed by transferring the samples from the gel to PVDF membrane. After blocking with 0.5 % blocking regent (Roche, 1096176)/PBS for 1 hr at room temperature (RT), the samples were immunoblotted for 1 hr at RT with anti-c-Kit, anti-EGFP (Clontech, 632569) and anti-β-actin (Santa Cruz, sc-4778) antibodies in 0.5% blocking regent/PBS at dilution of 1:500, 1:5000 and 1:2500, respectively. Following three times washing in 0.1% Tween-20 in Tris buffered saline (TBST, TBS: 140 mM NaCl, 2.7 mM KCl, 25 mM Tris, pH 7.4) for 5 min each at RT, the samples were incubated for 1 hr at RT with horseradish peroxidase-conjugated anti-rabbit IgG (GenScript, A00098) or anti-mouse IgG (Pronteintech, SA00001-1) antibodies at a dilution of 1:5000 and 1:10000, respectively. After three times washing in TBST, signals were detected using Chemi-Lumi One L (nacalai tesque, 07880) by the luminous image analyzer LAS-3000 mini (Fuji Film).

### Immunocytochemistry

Transfected DF1 were fixed in acetic acid/ethanol (1:5) for 10 min at RT, and washed in PBS three times for 5 min each at RT. The specimens were permeabilized in 0.1% Tween-20 in PBS for 10 min at RT, followed by washing in PBS twice for 5 min each at RT. After blocking with 0.5 % blocking reagent for 90 min at RT, the specimens were incubated overnight at 4 °C with dilution of 1:500 both anti-c-Kit and EGFP antibodies in 0.5% blocking regent/PBS. Following three times washing in PBS for 5 min each at RT, the specimens were incubated for 1 hr at RT with anti-rabbit IgG(H+L)-Alexa 568-conjugated antibody (Donkey; Invitrogen, A10042), anti-mouse IgG_2a_-Alexa 488-conjugated antibody (Goat; Invitrogen, A21131), and DAPI at a dilution of 1:500, 1:500 and 1:2000, respectively. After washing three times in PBS for 5 min at RT, fluorescent images were obtained using the Nikon A1R confocal microscope.

### Immunohistochemistry

A hindgut was dissected from chicken embryo, and embedded in FSC 22 Clear frozen section compound (Leica, 3801480). Cryostat sections of 10 μm were prepared (Thermo Scientific, Cryostar NX70). Following drying by cool air, the sections were fixed in acetic acid/Ethanol (1:5) for 10 minutes at RT. After washing in PBS for 10 minutes, the sections were incubated in 0.1% Tween in PBS for 10 minutes and washed by PBS. Subsequently, they were incubated with 1% blocking reagent/PBS for 1 hr at RT, followed by primary antibodies; 1:300 dilution of anti-c-Kit, 1:500 dilution of anti-αSMA (Sigma-Aldrich, A5228), 1:500 dilution of Tuj1 (Proteintech, 66375-1-Ig) overnight at 4 °C. After washing three times for 10 minutes each at RT, they were incubated with 1:500 dilution of anti-rabbit IgG(H+L)-Alexa 488-conjugated antibody, anti-mouse IgG_2a_ or IgG1-Alexa 568-conjugated antibody (Goat; Invitrogen, A21124), 1:1000 dilution of DAPI for 1.5 hrs at RT. After washing three times for 10 minutes each at RT, the specimens were sealed with Fluoromount (Diagnostic BioSystems). Fluorescent images were obtained using the Zeiss MZ10F microscope with Axiocam 503 mono camera.

### Whole-mount immunohistochemistry with tissue clearing

Hindgut and gizzard were dissected from chicken embryo and fixed in acetic acid/ethanol (1:5) overnight at 4 °C. The tissue clearing was conducted with a sorbitol-based optical technique as described previously (Hama et al., 2015). Briefly, specimens were incubated in a sorbitol-based solution for adaptation, and in urea-based solutions for permeabilization and clearing. After washing in PBS for 6 hrs, the specimens were incubated for 36-40 hrs at RT with 1:200 dilution of anti-c-Kit antibody, 1:300 dilution of anti-αSMA antibody in Absca*l*e solution, a PBS solution containing 0.33M urea and 0.5% TritonX-100. After removing primary antibodies, the tissues were washed in Absca*l*e solution twice for 2 hrs each at RT. Subsequently, they were incubated with 1:300 dilution of anti-rabbit IgG(H+L)-Alexa 488-conjugated antibody, anti-mouse IgG_2a_-Alexa 568-conjugated antibody, 1:1000 dilution of DAPI in Absca*l*e solution for 36-40 hrs at RT. After removing second antibodies, they were rinsed in Absca*l*e solution for 2 hrs and 0.05 % (w/v) Tween-20 in PBS containing 2.5% FBS twice for 2 hrs each at RT. The specimens were re-fixed in 4 % (w/v) PFA/ PBS for 1 hr and rinsed in PBS. Finally, the specimens were incubated in a Sca*l*e4 solution containing 40 % (w/v) sorbitol, 4 M urea for clearing overnight at RT. Fluorescent images were obtained using a confocal microscope (Nikon, A1R).

### Probe preparation for in situ hybridization

Chicken *c-Kit* cDNA was obtained using RNA extract of an E10 midgut with the primer set: forward 5-AGA CTA GTG TGC CCA CTA ACA GAT CCAG-3, reverse 5-GAC TCG AGT TAA TCG GGT AAG GTG AAG-3), and cloned into the pBluescript II SK plasmid (Stratagene). Digoxigenin-labeled RNA probe was prepared according to the manufacturer’s instruction (Roche).

### Whole-mount *in situ* hybridization

Whole mount in situ hybridization with a chicken gizzard was performed largely as previously described (Chevalier et al., 2020). A dissected E8 gizzard was fixed in 4 % PFA overnight at 4 °C, washed in PBST (0.1% Tween) twice for 5min each, and dehydrated through 25%, 50%, 75% and 100% methanol in PBST. When the specimens were stored at −20 °C, they were rehydrated in PBS and washed in PBST. The specimens were incubated for 1 hr in 6% hydrogen peroxide in PBST, and treated with proteinase K (10 μg/mL) for 10 min at RT, and washed with PBST. They were re-fixed in 0.2 % glutaraldehyde/4% PFA in PBST for 20 min at RT. After rinsing with PBST, specimens were prehybridized at 70 °C for 2 hrs, followed by hybridization with the probe (200 ng/ml) overnight at 70 °C. After washing at 65 °C, specimens were incubated in 20% FBS /PBST for 2.5 hrs at RT, and finally incubated with 1/2000 dilution of anti-DIG in 20% FBS/PBST overnight at 4 °C. The alkaline phosphatase activity was visualized by incubating the specimens in NTMT containing 0.07 mg/ml nitroblue-tetrazolium chloride (Roche), and 0.035 mg/ml 5-bromo-4-chloro-3-indolyphosphatase (Roche). Images were acquired by the Leica MZ10F microscope with the DS-Ri1 camera.

## Results and Discussion

### Polypeptides used for immunization

For the immunization of a rabbit, a polypeptide of 13 amino acids (aa) at the position of 716 to 728 of the chicken c-Kit protein was synthesized. Anti-serum harvested from an immunized rabbit was subjected to affinity purification using the aforementioned polypeptide. In the current study, the purified product is called a newly raised anti-c-Kit antibody (polyclonal).

### Validation of the newly raised c-Kit antibody by Western blotting assays

To evaluate the usefulness of the newly raised anti-c-Kit antibody by Western blotting, we prepared chicken fibroblast DF1 cells expressing EGFP, c-Kit, or c-Kit-EGFP (fusion) proteins (Fig. 1A). Since the Western blotting does not necessarily display a band at the predicted size of the protein, we started the analyses using a DF1 lysate containing the c-Kit-EGFP fusion protein, which must be detected by the anti-EGFP (Clontech, 632569) and newly raised anti-c-Kit antibodies. The predicted size of this fusion protein was 138 kDa. Multiple bands were detected at the size ranging from 130 kDa to 170 kDa by anti-EGFP antibody (Fig. 1B), which were also detected with the antic-Kit antibody (Fig. 1C). These bands were not seen in the lysate of EGFP/DF1 used as a control, whereas the EGFP protein in these cells was successfully detected by the anti-EGFP antibody (Fig. 1B, lanes “EGFP”). In the lysate of DF1 cells transfected with the *c-Kit* but not *EGFP* genes, the c-Kit antibody detected three bands of 109 kDa, 130kDa and 150kDa, which were not detected in EGFP/DF1 lysate (Fig. 1C). The band of 109 kDa matches the predicted size of the chicken c-Kit protein. These results suggest that the newly raised anti-c-Kit polyclonal antibody specifically detects the c-Kit proteins by the Western blotting assay.

The multiple sizes of the chicken c-Kit protein detected in Western blotting are consistent with those reported for the c-Kit protein in mice, showing 106 kDa, 124 kDa, and 160 kDa as modified products by glycosylation during transportation from endoplasmic reticulum (ER) to the plasma membrane via Golgi apparatus (Reith et al., 1991; Schmidt-Arras et al., 2005; Shi et al., 2016; Xiang et al., 2007).

### Validation of the newly raised c-Kit antibody by immunocytochemistry

We asked if the newly raised anti-c-Kit antibody was also available for immunocytochemistry. Since the widely used paraformaldehyde (PFA) fixative failed to yield satisfactory signals, we tested a variety of conditions to optimize the fixative. In the following studies, we used fixative of acetic acid: ethanol (1:5). DF1 cells expressing either the c-Kit-EGFP fusion protein or EGFP (control) were co-incubated with anti-EGFP antibody and the newly raised anti-c-Kit antibody.

DF1 cells expressing the c-Kit-EGFP fusion protein were successfully stained both by anti-EGFP and anti-c-Kit antibodies (Fig. 2A), whereas anti-c-Kit antibody did not yield signals in EGFP (control)-expressing DF1 cells (Fig. 2B). In magnified view, signals visualized by anti-c-Kit antibody were localized at the plasma membrane, which overlapped with signals stained by anti-EGFP antibody. Intercellularly, signals were also detected in the vicinity of the nucleus most likely in ER and/or Golgi apparatus. This notion is consistent with that immature c-Kit protein undergoes glycosylation in these intercellular components (Reith et al., 1991; Schmidt-Arras et al., 2005; Shi et al., 2016; Xiang et al., 2007). Of note, c-Kit-EGFP-expressing DF1 cells often exhibited multiple cellular processes which never occurs in control DF1 cells (Fig. 2B). It has yet to be studied if this is related to the multipolarity seen in c-Kit-positive melanocytes (Tadokoro et al., 2016) and ICCs (see below).

**Figure 2.**
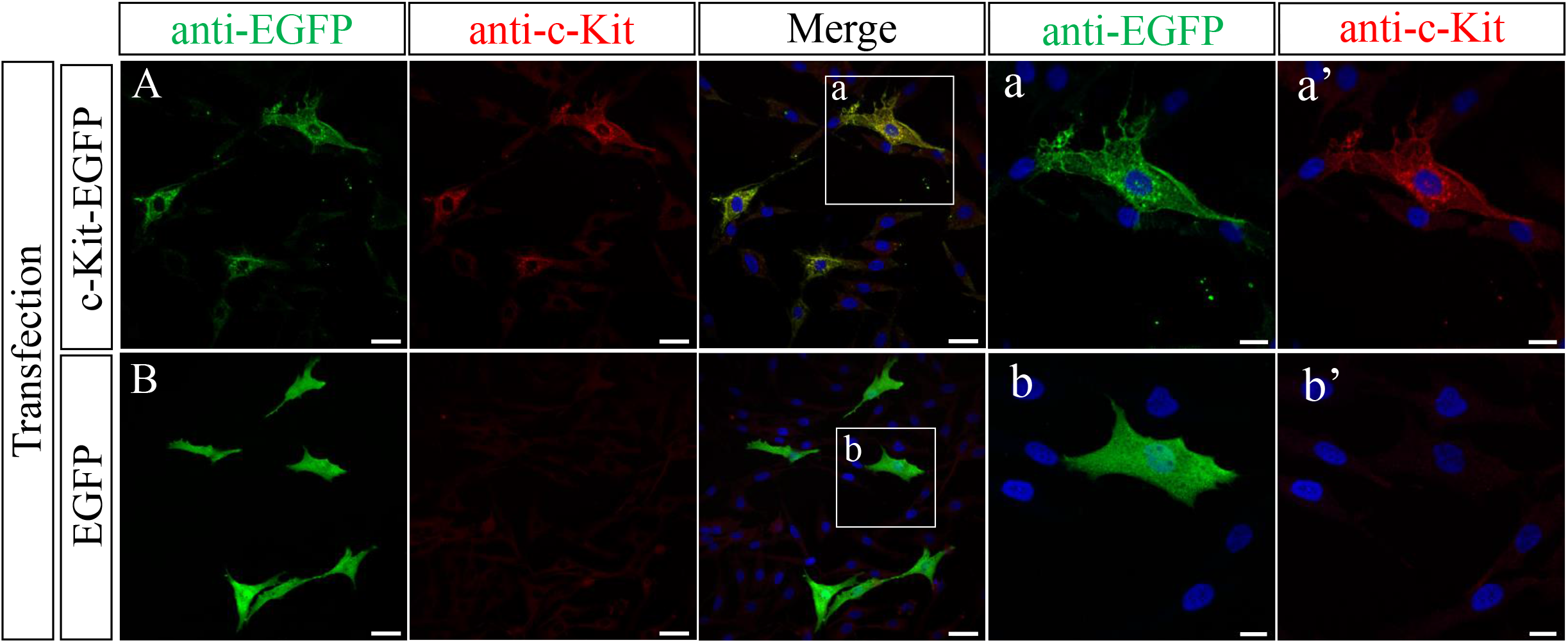
Immunostaining of c-Kit-EGFP/DF1 (A) and EGFP/DF1 (B) cells with the newly raised antibody and anti-EGFP antibody. (a, b) Magnified views of the square in (A, B). Scale bars: (A, B) 10 μm, (a, b) 20 μm.

### Validation of the newly raised c-Kit antibody to detect native proteins

To know if the newly raised c-Kit antibody can be used to examine the distribution of the native protein, we prepared a chicken gizzard at embryonic day 8 (E8), and compared patterns of signals detected by immunohistochemistry and *in situ* hybridization with the *c-Kit* mRNA as a probe. It was previously reported using whole mount in situ hybridization that the embryonic gizzard shows a characteristic pattern of *c-Kit* expression, a positive peripheral region enclosing an internal negative area (Chevalier et al., 2020). We indeed observed the identical pattern by *in situ* hybridization with the *c-Kit* mRNA (Fig. 3A). Importantly, this pattern was successfully recapitulated by whole mount immunostaining using the newly raised c-Kit antibody (Fig. 3C). At higher magnification, cluster-like patterns recognized by *in situ* hybridization were also detected by the anti-c-Kit antibody (Fig.3a, c). Taken together, we conclude that the newly raised anti-c-Kit antibody can be used for the detection of the native c-Kit protein in chicken embryos.

**Figure 3.**
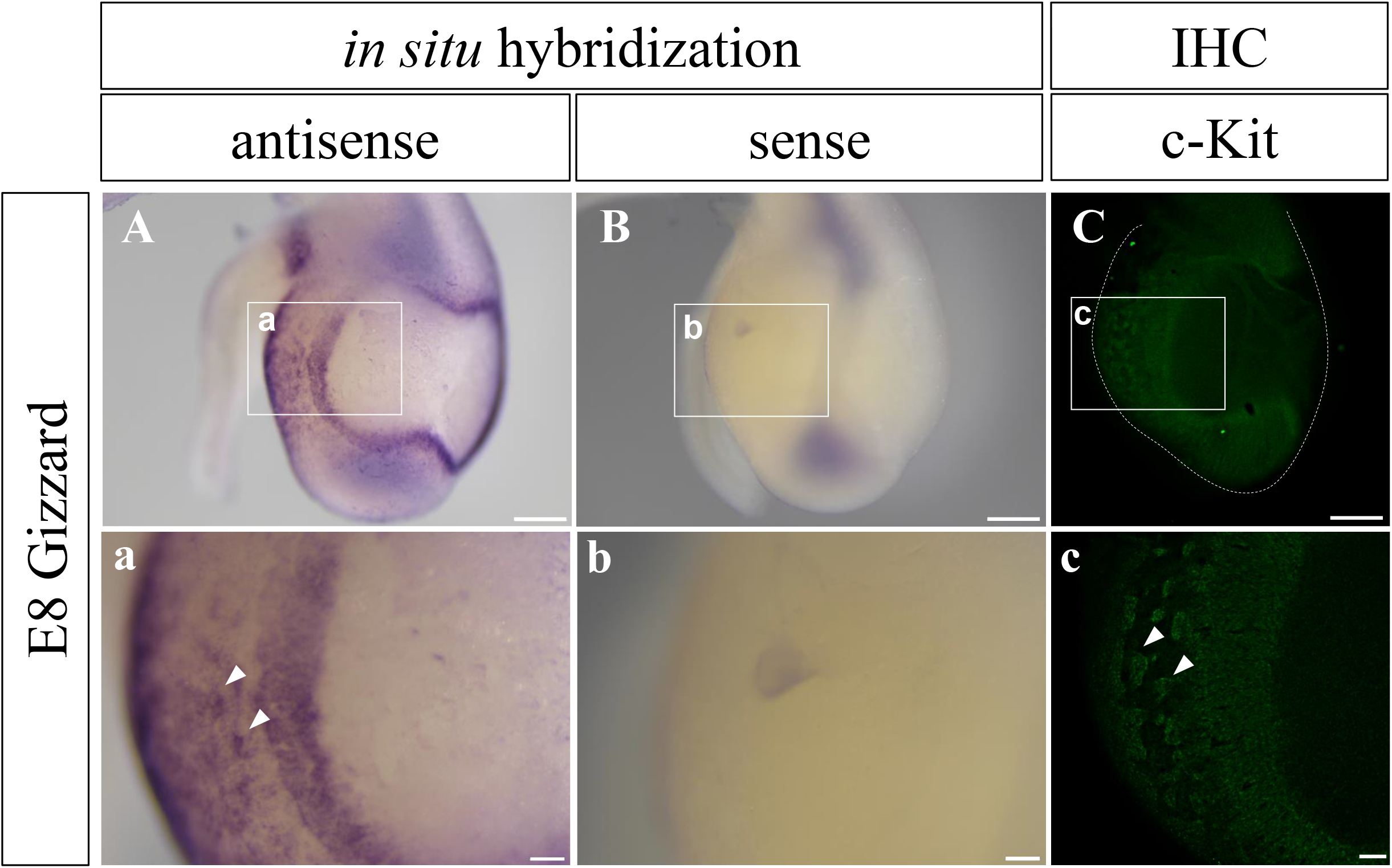
c-Kit signal in the gizzard at E8 was visualized by *in situ* hybridization with *c-Kit* probe (A, B), and whole-mount immunostaining with the new antibody (C). Signals patterns are almost identical. (a-c) Magnified views of the squares in (A-C). Arrowheads show cluster-like signals in (a, c). Scale bars: 500 μm in A-C, and 100 μm in a-c.

### ICCs were detected near the submucosal plexus in the embryonic hindgut

It is known in mice that common progenitors (naive cells) of ICCs and smooth muscles cells express both c-Kit and αSMA, whereas differentiated ICCs and smooth muscle cells are single positive for c-Kit and αSMA, respectively (Midrio et al., 2004; Radenkovic et al., 2010). We therefore co-stained transverse sections of embryonic hindgut with anti c-Kit antibody and commercial antibody for αSMA (sigma) to detect differentiating ICCs and smooth muscles. Hindguts dissected from embryos at stages ranging from E7 to E18 were fixed in acetic acid: ethanol (1:5).

In the hindgut at E7 and E9, c-Kit signals were uniformly distributed in the CM layer, and this pattern overlapped with that of αSMA (Fig.4B-E; not shown for E9), implying that the majority of the gut mesodermal cells are common progenitors for ICCs and SM cells at these stages (LM is formed at E11 onward (Huycke et al., 2019)). By E15, c-Kit single positive cells emerged. A prominent example was the signal observed at E18 in the layer of submucosa which lies in between the CM layer and muscularis mucosa (Fig. 4F-I). Along the layer of muscularis mucosa, three sub-layers were recognized in a concentric manner: αSMA^+^/c-Kit^−^, αSMA^+^/c-Kit^+^, and αSMA^−^/c-Kit^+^ in a central to peripheral direction (Fig. 4A, F-I). It is possible that common progenitors (αSMA^+^/c-Kit^+^) have split to yield the smooth muscle cells (αSMA^+^/c-Kit^−^) centrally, and the ICCs (αSMA^−^/c-Kit^+^) peripherally. These ICCs correspond to ICC submucosa (ICC-SM) reported in adult chicken (Yang et al., 2012). Along a radial direction, a stream-like pattern of αSMA^−^/c-Kit^+^ population (ICCs) was observed that extended from the submucosal layer to the CM layer (Fig. 4A, F-I). These ICCs might transmit signals between the submucosal and CM layers. We further noticed that this extension of ICCs was frequently adjacent to submucosal plexus of ENS visualized by Tuj1 (Fig. 4A, J-M), suggestive of signal communications between ICC-SM and submucosal plexus (ENS). The radially extended ICC population is unprecedented in mammals, and it has yet to be studied if these cells are specific to chickens. Together, co-staining with antibodies including our new antibody has revealed novel patterns of ICCs in the submucosal layer. At the level of the resolution of this immunohistochemistry, it was not clear whether differentiated ICCs (c-Kit single positive) were present in the CM layer since the majority of the CM cells were faintly stained by both anti-c-Kit and αSMA (see below).

**Figure 4.**
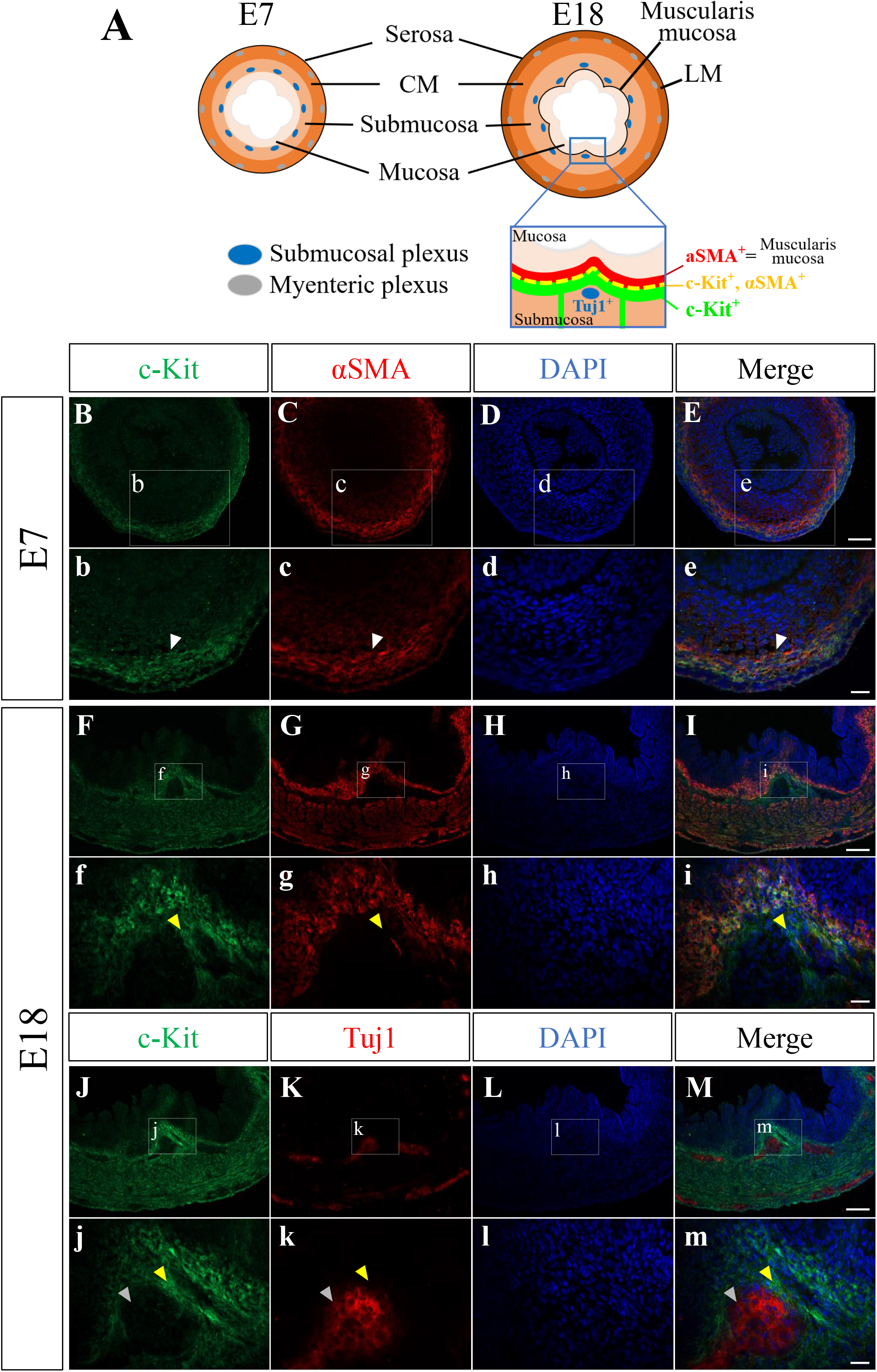
Immunohistochemical assays with cryo-sections of embryonic hindgut. (A) Schematic drawings show concentric layered structures of the gut at E7 and E18. The magnified square in E18 corresponds to the views in I, M. (B-E) Staining of E7 hindgut with anti-c-Kit and αSMA antibodies. (F-I) Co-staining of E18 with anti-c-Kit and αSMA antibodies. (J-M) Co-staining of E18 with anti-c-Kit and Tuj1 antibodies, the latter being a marker for differentiating neuron-specific class III-β tubulin. (b-m) Magnified views of the square in (B-M). White arrowhead shows co-expression of c-Kit and αSMA in (B, C, E). Yellow arrowhead shows the extension of cell population of c-Kit^+^/αSMA^−^ along the radial axis, located adjacent to Tuj1-positive submucosal plexus (grey arrowhead). Scale bars: 50 μm in B-E, 100 μm in F-M, 20 μm in b-m.

### The newly raised antibody visualized individual ICCs in a cleared hindgut

To further scrutinize the morphology of individual ICCs in the developing hindgut, the whole mount immunohistochemistry was combined with the tissue clearing technique, which facilitates visualization of fluorescent signals at high resolution (Hama et al., 2015; Watanabe et al., 2017) (also see Material and Methods). A piece of E18 hindgut was costained with the new anti-c-Kit antibody and anti-SMA antibody, and subjected to the tissue clearing process. By confocal microscopy, c-Kit single positive ICCs (green) were observed in the CM layer, and they were bipolar in shape and aligned with elongated muscle cells (red) (Fig. 5A, B). In contrast, ICCs residing in a layer between the CM and LM (corresponding to the layer of myenteric plexus) showed multipolarity, and were not aligned with smooth muscles (Fig. 5C, D). ICCs were also found in the LM layer, and they were not bipolar (Fig.5E, F). Although the immunohistochemistry by our new antibody failed to delineate a cell shape, cell morphology/shape of ICCs could be deduced by their relative positions with neighboring cells and also by a shape of a nucleus visualized by DAPI staining in a range of z-stack images of confocal microscopy. Unlike mice, ICCs in the hindgut did not form a prominent network by E18 (Torihashi et al., 1997).

**Fig. 5.**
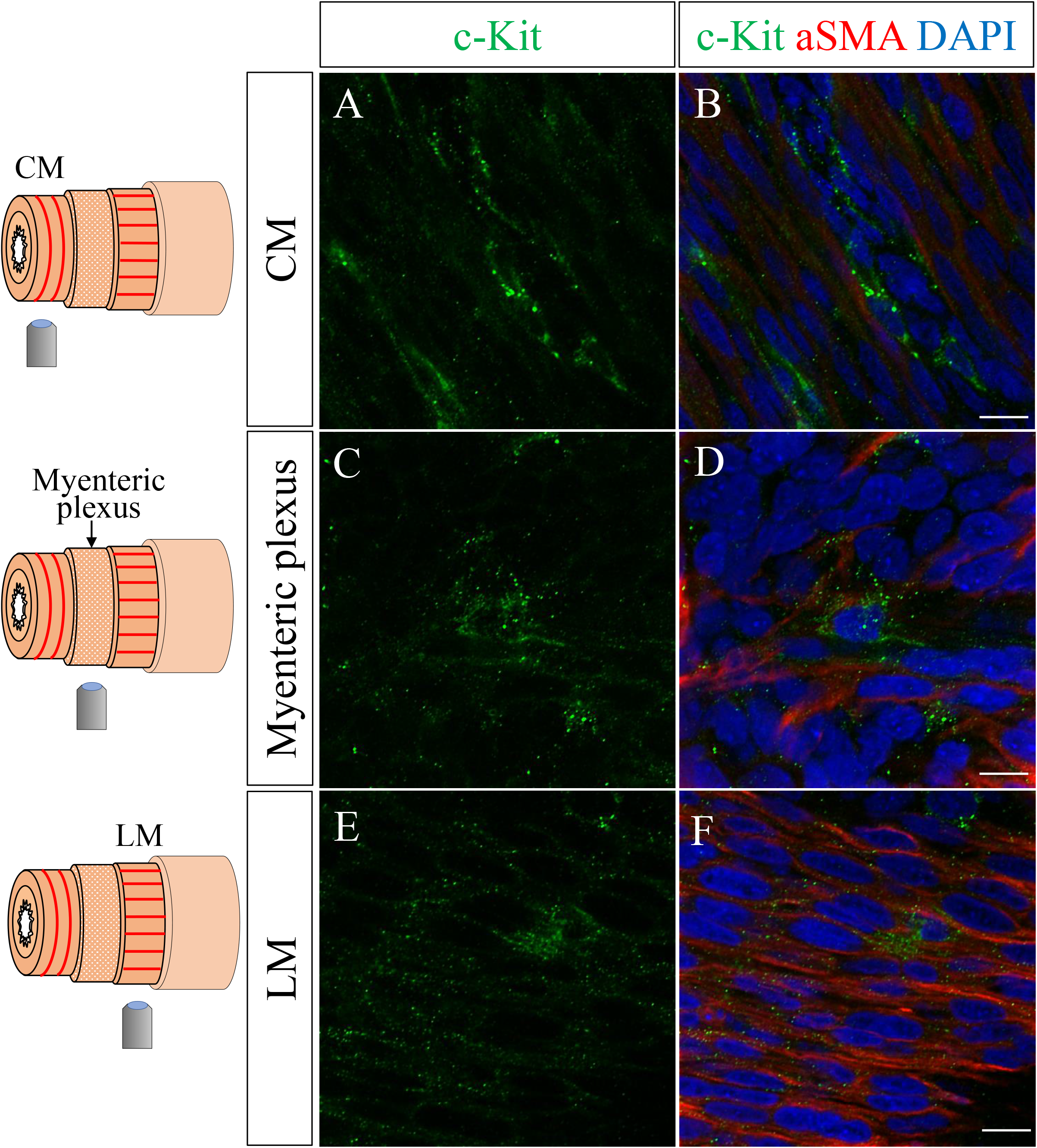
Co-staining with anti-cKit and αSMA antibodies visualized individual ICCs in cleared hindgut at E18. Three different focal layers of the hind gut, identified by characteristic alignments of smooth muscle cells (red lines), were assessed by confocal microscopy. CM (A, B), myenteric plexus (C, D), and LM (E, F). Scale bars: 10 μm.

In summary, the polyclonal antibody obtained in this study successfully recognizes and visualizes the chicken c-Kit protein in embryonic tissues, and therefore this polyclonal antibody is useful to investigate differentiating ICCs and ICCs’ contribution to the coordinated peristaltic movements in the developing gut.

## Acknowledgement

This work was supported by JSPS KAKENHI Grant Numbers; 20K21425, 20H03259, 19H04775, and also by Research Foundation for Opto-Science and Technology. R. Y. and Y.S are a fellow and ex-fellow of JSPS, respectively.

